# Ancient symbiosis confers desiccation resistance to stored grain pest beetles

**DOI:** 10.1101/182725

**Authors:** Tobias Engl, Nadia Eberl, Carla Gorse, Theresa Krüger, Thorsten H. P. Schmidt, Rudy Plarre, Cornel Adler, Martin Kaltenpoth

## Abstract

Microbial symbionts of insects provide a range of ecological traits to their hosts that are beneficial in the context of biotic interactions. However, little is known about insect symbiont-mediated adaptation to the abiotic environment, e.g. temperature and humidity. Here we report on an ancient (~400 Mya) clade of intracellular, bacteriome-located Bacteroidetes symbionts that are associated withgrain and wood pest beetles of the phylogenetically distant families Silvanidae and Bostrichidae. In the saw-toothed grain beetle Oryzaephilus surinamensis, we demonstrate that the symbionts affect cuticle thickness, melanization and hydrocarbon profile, enhancing desiccation resistance and thereby strongly improving fitness under dry conditions. Together with earlier observations on symbiont contributions to cuticle biosynthesis in weevils, our findings indicate that convergent acquisitions of bacterial mutualists represented key adaptations enabling diverse pest beetle groups to survive and proliferate under the low ambient humidities that characterize dry grain storage facilities.

## Introduction

Microbial mutualists are a major driving force of evolution (Klepzig et al., 2009), as they confer a variety of ecological benefits to their host (Feldhaar, 2011, Oliver and Martinez, 2014). In insects, numerous studies yielded evidence for symbiont-provided benefits in the context of biotic interactions, particularly through the supplementation, degradation, or detoxification of the diet (Moran, 2006, Douglas, 2009, van den Bosch and Welte, 2017) or by defending the host against natural enemies (Florez et al., 2015). However, comparatively little is known about symbiont-mediated adaptations to the abiotic environment. Notably, several studies reported on symbionts that enhance resistance of insects to high temperatures (Russell and Moran, 2006, Montllor et al., 2002, Brumin et al., 2011). In most of these cases, however, facultative secondary symbionts ameliorate heat susceptibility of primary obligate symbionts rather than directly altering the host physiology and thereby extending the viable temperature range (Wernegreen, 2012, Corbin et al., 2017). Nevertheless, under selective conditions for heat resistance, during hot summers in desert sites, the abundance of protective, secondary symbionts was indeed found to increase in pea aphids, presumably reflecting adaptation to higher temperatures (Harmon et al., 2009).

Stored grain pest insects profit from an excess of food but face the challenge of low environmental humidity that is maintained in storage facilities to prevent the growth of mould fungi (Hagstrum et al., 1996). Several groups of beetles independently managed to invade the same ecological niche of stored grain and dried plant products despite the considerably lower humidity compared to the ancestral habitat associated with a fungivorous or saprophagous state of living under bark (Crowson, 1981, Hunt et al., 2007). Most of these groups were described to harbor facultative intracellular bacterial symbionts. Weevils of the genus *Sitophilus* engage in a symbiosis with the γ-proteobacterium *Sodalis pierantonius*, and the silvanid saw-toothed grain beetle *Oryzaephilus surinamensis* and several bostrichid beetles are associated with as yet unidentified symbionts (Koch, 1931, Mansour, 1934, Koch, 1936a, Buchner, 1965, Nardon and Grenier, 1988, Heddi et al., 1999, Kleespies et al., 2001). In all cases, the symbionts can be experimentally depleted or removed without disrupting the hosts’ life cycle in laboratory settings, and some populations of *O. surinamensis* were even found to contain aposymbiotic individuals in the field (Koch, 1936b, Huger, 1956, Vigneron et al., 2014). However, *S. pierantonius* was shown to provide precursors for cuticle biosynthesis that are especially important during beetle development, and symbiont cells are actively degraded in adults (Vigneron et al., 2014). Symbiont-free (aposymbiotic) weevil populations suffer from a lower growth rate due to higher mortality and reduced fecundity, but are viable and able to reproduce (Nardon and Grenier, 1988). The symbionts of both *O. surinamensis* and *Rhizopertha dominica* are likewise not obligate, and past studies were unable to establish any evidence for a physiological or ecological benefit of their presence (Koch, 1956, Huger, 1956).

Here, we investigate the hypothesis that engaging in mutualistic associations with bacteria represents a (pre)adaptation in several beetle families to exploit stored grain products as a food source. To that end, we characterized the intracellular symbionts of five bostrichid and three silvanid species of grain and wood pest beetles, revealing a shared and ancient symbiosis related to the intracellular Bacteriodetes symbionts of cicadas *(Sulcia)*, cockroaches and termites *(Blattabacterium).* For the saw-toothed grain beetle *O.surinamensis*, experimental symbiont elimination resulted in reduced cuticle melanization and thickness as well as increased cuticular hydrocarbon biosynthesis upon drought stress. Concordantly, aposymbiotic beetles suffered from lower population growth rates and were more susceptible to desiccation and drought-inflicted mortality, indicating that the symbiosis enhances desiccation resistance and thereby likely played, besides in other important functions, like the defense against natural enemies, a key role for the adaptation of phylogenetically diverse beetles to conditions of low ambient humidity in mature grain and especially stored grain facilities.

## Results

### Bostrichid and silvanid pest beetles harbor ancient *Sulcia-like* intracellular symbionts

By PCR amplification, cloning and sequencing of the bacterial 16S rRNA gene, we characterized the symbionts associated with five bostrichid beetles (*Lyctus brunneus, Rhizopertha dominica, Prostephanus truncatus, Dinoderus bifoveolatus* and *Dinoderus porcellus*) and three silvanid beetles *(Ahasverus advena, Oryzaephilus mercator* and *Oryzaephilus surinamensis*). While *L. brunneus* feeds on seasoned hard wood, all others are serious pests of diverse stored grain products. Surprisingly, despite the phylogenetic distance of Bostrichidae and Silvanidae (about 240 mya, see Hunt et al., 2007), the symbionts of all eight species were assigned to the same clade of Bacteroidetes bacteria that also contained *Sulcia muelleri*, the symbionts of Auchenorrhyncha, and *Blattabacterium*, the symbionts of cockroaches and some termites (Fig. 1, Supplemental Fig. 1). While the three silvanid species and two of the bostrichids *(R. dominica* and *P. truncatus)* contained a single symbiont, *L. brunneus* and the two *Dinoderus* species additionally displayed a second, more basally branching clade of symbionts (Fig. 1). Based on diagnostic PCRs, the derived and in all species maintained symbiont could be detected in 68%-99% (Supplemental Table 2) of tested adult beetles except *A. advena*, whereas the ancestral symbiont only associated with *Lyctus* and *Dinoderus* was detected in 68% −90%. In total, 95%-100% of the tested individuals were positive for at least one of both symbionts. In *O. surinamensis* and *R. dominica*, the degradation of symbionts in old individuals (particularly in males) has been reported (Huger, 1956), probably accounting for the low apparent infection frequencies across host species. Infection rates estimated by diagnostic PCR and Fluorescence in situ hybridization (FISH) in *A. advena* were with 30% considerably lower (N=10). Due to rare and low levels of infections (Fig. 2d) symbionts in *A. advena* were probably formerly not detected (Buchner, 1965). Consistenlty, despite being usually also considered as a stored grain pest, *A.advena* actually feeds on fungal infestations of grain, requires the addition of yeast extract in artificial grain diets and also relatively high environmental humidities of 70% (Thomas and Leschen, 2009).

**Figure 1.**
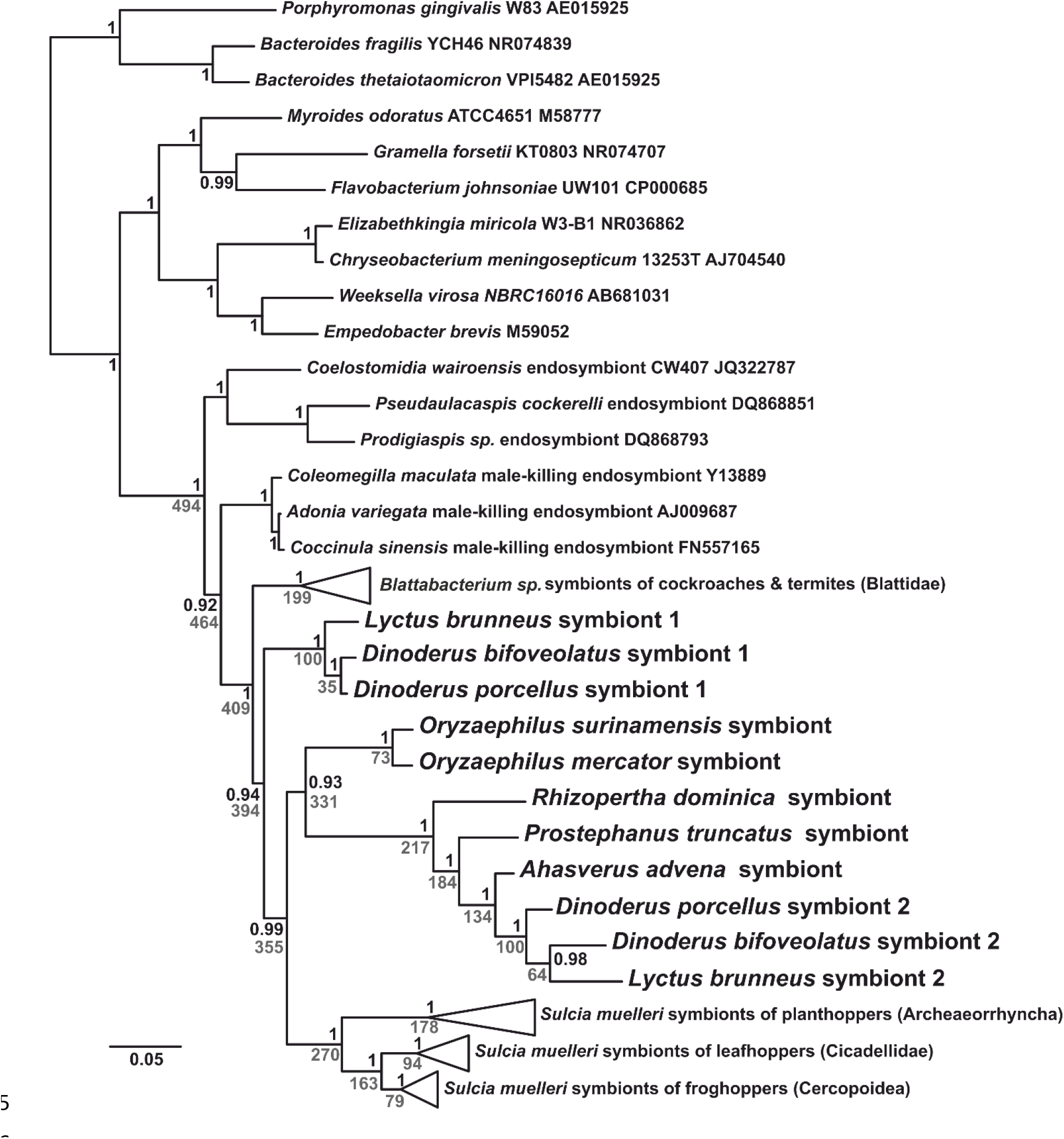
Phylogenetic placement of intracellular symbionts in silvanid and bostrichid grain pest beetles within the Bacteroidetes, and their close association to endosymbionts of cockroaches and cicadas. The phylogeny was reconstructed using Bayesian inference, and black node values represent Bayesian posteriors. Grey values represent mean node ages in Mya, based on a phylogenetic dating analysis with the age of *Blattabacterium* and *Sulcia* as calibration points.

**Figure 2.**
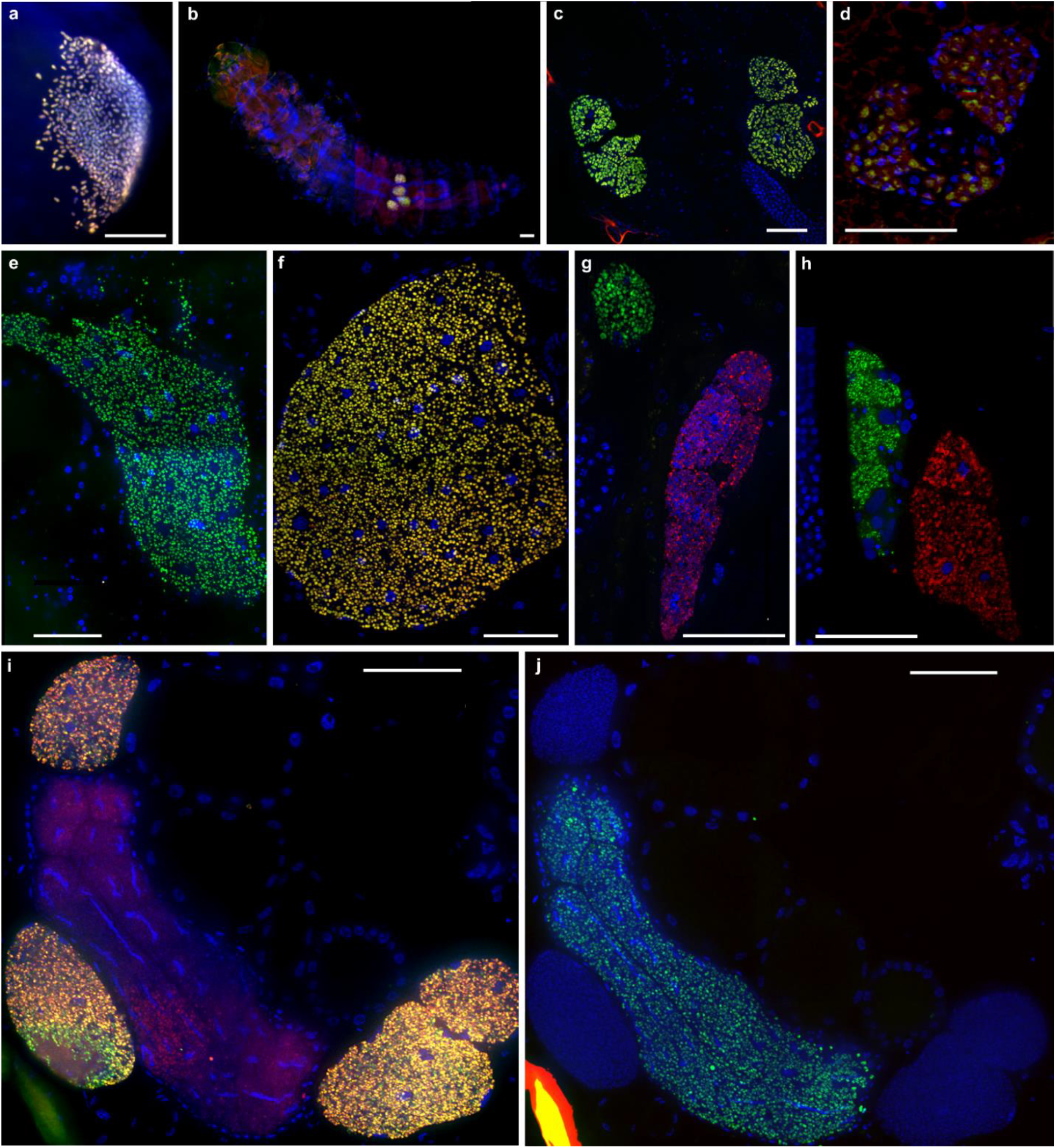
Bacteroidetes symbiont localization in silvanid and bostrichid beetle bacteriomes. Whole mount Fluorescence *in situ* hybridization (FISH) of *O. surinamensis* (a) egg, (b) larva, and (c) a longitudinal section of an adult female stained with EUB338-Cy3 (red) and OsurSym16S-Cy5 (green). (d) Cross section of an *A. advena* adult stained with Eub338-Cy3 and CFB563-Cy5. Longitudinal sections of (e) *R. dominica* and (f) *P. truncatus* adults, stained with EUB338-Cy3 (red; did not work in e) and Bostrichidae_Sym2-Cy5 (green), of adult (f) *D. bifoveolatus* and (g) *D. porcellus*, stained with Bostrichidae_Sym1-Cy3 (red) and Bostrichidae_Sym2_Cy5 (green). Cross sections of a *L. brunneus* female stained with (i) Eub338-Cy3 (red) and Bostrichidae_Sym1-Cy5 (green) and (j) Bostrichidae_Sym2-Cy5 (green). DAPI was used as a general DNA stain for all experiments (blue). Scale bars represent 50μm.

A phylogenetic dating analysis based on the partial symbiont 16S rRNA gene of ~1250bp and two calibration points, the origin of the cicada-*Sulcia* symbiosis (260-280Mya; Moran et al., 2005) and the cockroach-*Blattabacterium* symbiosis (150-300Mya; Patino-Navarrete et al., 2013), revealed an ancient origin of the clade of Bacteroidetes endosymbionts 494 Mya ago (Fig 1, Supplemental Fig 1 and Supplemental Table 3). The mutualistic Bacteroidetes group comprising *Blattabacteria*, the beetle symbionts and *Sulcia* dates back around 394 Mya, with the split of the ancestral beetle symbiont clade and the split between *Sulcia* and the other beetle symbionts around 355Mya ago and the split of the *Oryzaephilus* and bostrichid symbionts around 331 Mya ago (Fig 1, Supplemental Fig 1 and Supplemental Table 3). Different nucleotide substitution models had little impact on node ages with a mean node age for all endosymbionts varying between 489-503 Mya, and the mutualistic symbionts between 404-414 Mya, whereas strict clock models resulted in younger node ages (423-432 Mya and 384-392 Mya, respectively, see Supplemental Table 3). Omitting one calibration point shifted the divergence times to a considerably earlier time of 593 Mya, if only the origin of *Blattabacterium* was included, and to 474 Mya with only the origin of *Sulcia* included (Supplemental Table 3).

The bacterial symbionts were all located intracellularly, in bacteriomes located between gut, fat body and reproductive organs, but without direct connection to any of these tissues. While the Silvanid beetles contained two pairs of bacteriomes, both with the same single symbiont strain (Fig. 2a-d; see also Koch, 1936a), *R. dominica* and *P. truncatus* contained only one bacteriome pair with a single strain (Fig. 2e+f). In contrast, both *Dinoderus* species contained one pair of bacteriomes for each symbiont strain, which were anatomically separated from each other (Fig. 2g+h). *L. brunneus* harbored a pair of bacteriomes, each composed of a central bacteriome containing the ancestral symbiont strain surrounded by three bacteriomes harboring the derived symbiont strain (Fig. 2i+j; see also Koch, 1936a).

### Symbiont association is not obligate for *O. surinamensis*

As described previously, the *O. surinamensis* symbiont titers could be reduced by either heat or tetracycline treatment (Koch, 1936b, Huger, 1956). Treatment of adult beetles over three months with tetracycline resulted in a complete and stable elimination of the symbiont, while heat treatment only reduced symbiont titers (Supplemental Fig. 2). In the tetracycline treatment group, the native symbiont could neither be detected by quantitative PCR (Supplemental Fig. 2a) nor by FISH (16 eggs and 8 adults tested per time point, 100% lacked a symbiont in the treatment group, whereas 100% in the control group showed infection). Although the qPCRs occasionally resulted in off-target amplification in the absence of the native symbiont, none of the amplification products in the tetracycline-treated group matched the melting profile of the native symbiont amplicon (Supplemental Fig. 2a). In contrast, the offspring (eggs) of beetles that were exposed to 36°C as either adults or larvae showed slightly, but not significantly lower symbiont titers (Mann-Whitney-U tests; heat treated larvae vs. control U=30, N=10 each, p=0.243, heat treated adults vs. controls U=101, N=30 and 10, p=0.269; Supplemental Fig. 2b). In the next generation, symbionts were significantly reduced and often completely absent (Mann-Whitney-U test, F2 of heat treated adults vs control, U=10.5, N=10 each, p=0.003; Supplemental Fig. 2b), but less consistently so than in the tetracyclin treatment group.

Due to the successful elimination of symbionts by tetracycline treatment, the offspring of tetracycline treated beetles were maintained in a continuous laboratory culture over two years to perform all following experiments. Despite overall good performance of aposymbiotic cultures under optimal growth conditions of 60% RH at 30°C, we observed a significant reduction of cuticle melanization (exact 2-sided Mann-Whitney-U test, U=154, p<0.001, Supplemental Fig. 3a) and a significantly lower population growth of aposymbiotic beetles (exact 2-sided Mann-Whitney-U test, U=30, p=0.004, Supplemental Fig. 3b).

### Symbionts contribute to cuticle thickness and melanization in *O. surinamensis*

Melanization of the cuticle increases its physical strength and contributes to desiccation resistance (Gibbs and Rajpurohit, 2010). Given the observed impact of symbiont elimination on cuticle melanization, we set out to assess the contribution of the *O. surinamensis* symbiont to cuticle formation and melanization in more detail, by exposing replicate symbiotic and aposymbiotic populations to dry (30-40%RH) and humid (60% RH) conditions. In adult beetles, both symbiont absence and reduced environmental humidity significantly reduced cuticle melanization (Table 1, Fig. 3a) and cuticle thickness (Table 1, Fig. 3b) with both a thinner endo- and exocuticle (Table 1, Supplemental Fig. 4a+b). Aposymbiotic adults exhibited overall a less melanized and on average 26% thinner cuticle than their symbiotic counterparts. In 4^th^ instar larvae, symbiont absence, but not the humidity regime, resulted in a significant reduction in cuticle thickness by about 20% (Table 1, Supplemental Fig. 4c).

**Figure 3.**
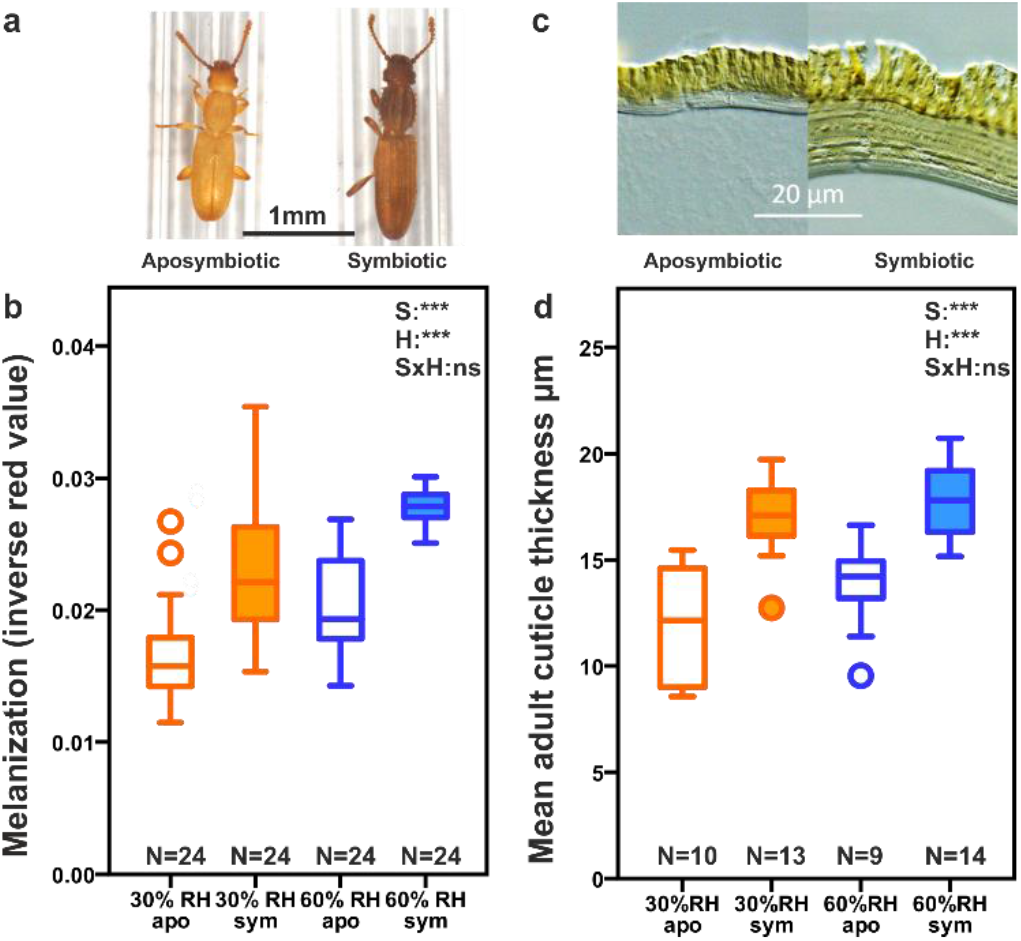
Melanization and cuticle thickness of symbiotic and aposymbiotic *O. surinamensis* adults. (a) Photographs of 2 day old aposymbiotic and symbiotic *O.surinamensis* adults and (b) melanization measured as thorax coloration of aposymbiotic and symbiotic adults reared at different humidities. (c) Ventral, thoracal cuticle sections of aposymbiotic and symbiotic *O.surinamensis* adults and (d) cuticle thickess of aposymbiotic and symbiotic adults reared at different humidities. Both symbiont presence (S) and environmental humidity (H; GLM, p<0.001), but not their interaction (S*H; GLM, p>0.05) had a highly significant influence on cuticle melanization and thickness. Boxplots show medians, quartiles and minima/maxima. Sample size is given under each box. Filled boxes represent symbiotic and empty ones aposymbiotic beetles, orange boxes indicate rearing at 30% RH, blue ones at 60% RH

**Table 1.** Statistical test results of generalized linear and cox-mixed effects models describing the influence of symbiont presence/absence, environmental humidity and acute desiccation stress on variable beetle parameters. Tests are in order as they appear in the manuscript. χ^2^ = χ^2^ Wald factors significantly influencing a parameter are highlighted in bold.

|  | test | symbiont presence/absence | Environmental humidity | acute desiccation stress | symbiont* humidity | symbiont* stress | humidity* stress | all three factors |
| --- | --- | --- | --- | --- | --- | --- | --- | --- |
| Melanization | GLM | $\chi^2=92.3$,<br>d.f.=1,<br>**p<0.001** | $\chi^2=94.1$,<br>d.f.=1,<br>**p<0.001** | | $\chi^2=0.5$,<br>d.f.=1,<br>p=0.488 | | | |
| Cuticle thickness | GLM | $\chi^2=52.5$,<br>d.f.=1,<br>**p<0.001** | $\chi^2=4.5$,<br>d.f.=1,<br>**p=0.034** | | $\chi^2=0.837$,<br>d.f.=1,<br>p=0.360 | | | |
| Endocuticle thickness | GLM | $\chi^2=32.6$,<br>d.f.=1,<br>**p<0.001** | $\chi^2=6.8$,<br>d.f.=1,<br>**p=0.009** | | $\chi^2=1.7$,<br>d.f.=1,<br>p=0.197 | | | |
| Exocuticle thickness | GLM | $\chi^2=49.5$,<br>d.f.=1,<br>**p<0.001** | $\chi^2=0.07$,<br>d.f.=1,<br>p=0.790 | | $\chi^2=0.04$,<br>d.f.=1,<br>p=0.843 | | | |
| Larval cuticle thickness | GLM | $\chi^2=17.8$,<br>d.f.=1,<br>**p<0.001** | $\chi^2=0.2$,<br>d.f.=1,<br>p=0.621 | | $\chi^2=0.5$,<br>d.f.=1,<br>p=0.473 | | | |
| Total CHC amounts | GLM | $\chi^2=14.6$,<br>d.f.=1,<br>**p<0.001** | $\chi^2=83.5$,<br>d.f.=1,<br>**p<0.001** | $\chi^2=3.5$,<br>d.f.=1,<br>p=0.061 | $\chi^2=6.2$,<br>d.f.=1,<br>**p=0.013** | $\chi^2=9.2$,<br>d.f.=1,<br>**p=0.002** | $\chi^2=0.7$,<br>d.f.=1,<br>p=0.391 | $\chi^2=5.4$,<br>d.f.=1,<br>**p=0.02** |
| Proportion unsaturated CHCs | GLM | $\chi^2=14.6$,<br>d.f.=1,<br>**p<0.001** | $\chi^2=83.5$,<br>d.f.=1,<br>**p<0.001** | $\chi^2=4.8$,<br>d.f.=1,<br>**p=0.029** | $\chi^2=1.9$,<br>d.f.=1,<br>p=0.169 | $\chi^2=0.7$,<br>d.f.=1,<br>p=0.418 | $\chi^2=5.9$,<br>d.f.=1,<br>p=0.015 | $\chi^2=0.8$,<br>d.f.=1,<br>p=0.366 |
| Average CHC chain length | GLM | $\chi^2=13.3$,<br>d.f.=1,<br>**p<0.001** | $\chi^2=0.02$,<br>d.f.=1,<br>p=0.903 | $\chi^2=0.9$,<br>d.f.=1,<br>p=0.356 | $\chi^2=7.1$,<br>d.f.=1,<br>**p=0.008** | $\chi^2=1.3$,<br>d.f.=1,<br>p=0.259 | $\chi^2=0.5$,<br>d.f.=1,<br>p=0.490 | $\chi^2=0.3$,<br>d.f.=1,<br>p=0.567 |
| Water loss | repeated measures GLM | $\chi^2=35.8$,<br>d.f.=1,<br>**p<0.001** | $\chi^2=217.2$,<br>d.f.=1,<br>**p<0.001** | | $\chi^2=0.6$,<br>d.f.=1,<br>p=0.449 | | | |
| Mortality | Cox mixed-effects model | $z=-2.11$,<br>**p=0.034** | $z=-1.22$,<br>p=0.220 | | $z=0.73$,<br>p=0.470 | | | |
| Population growth | GLM | $\chi^2=431.3$,<br>d.f.=1,<br>**p<0.001** | $\chi^2=926.5$,<br>d.f.=1,<br>**p<0.001** | | $\chi^2=74.1.1$,<br>d.f.=1,<br>**p<0.001** | | | |

### Effect of symbionts on the epicuticular hydrocarbon profile

By adapting their cuticular hydrocarbon (CHC) profile, insects including *O. surinamensis* can rapidly change the water permeability of their epicuticle (Howard et al., 1995, Gibbs and Rajpurohit, 2010). We exposed adult beetles from all four treatments (full factorial design of dry and moist, symbiotic and aposymbiotic) to severe desiccation stress (one day at <2% RH) or not (control) and measured their respective CHC profiles. Symbiont presence and environmental humidity during rearing had a significant influence on the total amount of CHCs, with symbiont elimination and dry conditions resulting in an increased amount of CHCs (Table 1). While acute drought stress alone did not affect CHC amounts (Table 1), the interaction with symbiont status did, with aposymbiotic beetles applying more CHCs to their cuticle under both chronic and acute desiccation stress (Table 1, Fig 4a). In addition, symbiont absence and long term exposure to low humidity significantly increased the proportion of unsaturated hydrocarbons (Table 1, p<0.001), and acute desiccation stress increased the proportion of unsaturated hydrocarbons in beetles adapted to low humidity, but decreased it for beetles adapted to high humidity (Table 1, Fig. 4b). Furthermore, symbiont absence itself and its interaction with environmental humidity also affected the average chain length of CHCs on the cuticle with aposymbiotic beetles carrying shorter chain CHCs, which is even enhanced under low humidity (Table 1, Fig. 4c), whereas neither long-term low humidity *per se* nor acute desiccation stress affected CHC chain length (Table 1, Fig 4c). These results demonstrate that aposymbiotic beetles perceive desiccation stress significantly more strongly than symbiotic beetles, especially if they were already kept under low humidity, and mount a physiological response to improve their epicuticular properties, as higher amounts of hydrocarbons provide a better evaporation protection. Shorter and less saturated hydrocarbons are generally considered to offer less protection against desiccation. However these conclusens are derived by studying the behavior of CHC mixtures at different temperatures, not at a fixed temperature with varying humidities (Gibbs and Rajpurohit, 2010). Accordingly, beetles without symbionts and under chronic, low humidity seem not to be able to keep the potentially more protective composition of their CHCs, but rather rely on the protective effect of a thicker hydrocarbon layer, whereas symbiotic beetles reared in a more humid environment are able to shift their CHC profile to a more favourable composition under acute desiccation stress.

**Figure 4.**
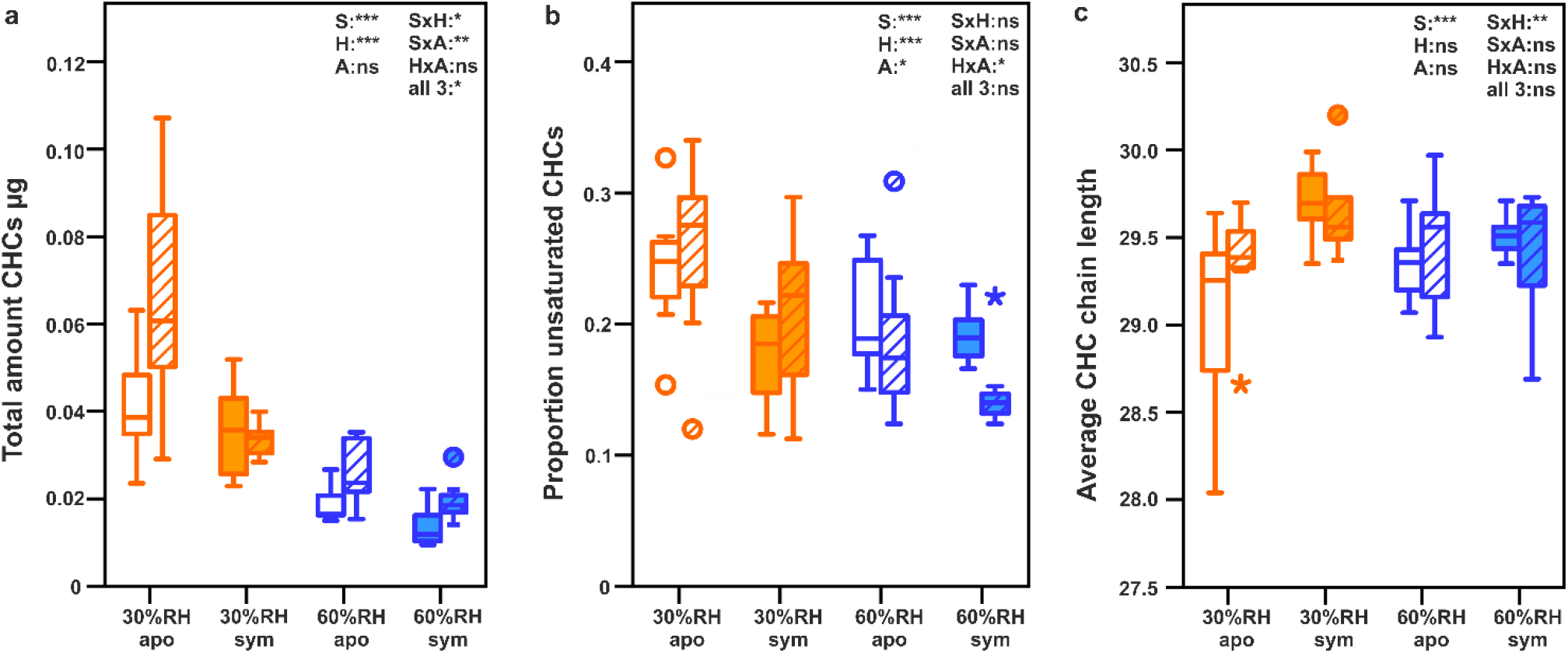
Rapid desiccation hardening (changes in the cuticular hydrocarbon profile) of *O. surinamensis* as a response to symbiont loss, environmental humidity and acute desiccation stress. (a) Total amounts of cuticular hydrocarbons per beetle (average calculated from 30 pooled beetle extracts), (b) proportion of unsaturated hydrocarbons and (c) average hydrocarbon chain length show physiological counteradaptations of beetles to long-term exposure to low environmental humidity, but especially to acute desiccation stress. Statistical results report on different factors and their interaction affecting CHC parameters (GLM, S=symbiont presence, H=environmental humidity, A=acute desiccation stress, ***: p<0.001; **: p<0.01; *: p<0.05, n.s.: p≥0.05). Filled boxes represent symbiotic and empty ones aposymbiotic beetles, orange boxes indicate rearing at 30% RH, blue ones at 60% RH, and hatched boxes show the respective changes after acute desiccation stress.

### Symbionts confer desiccation resistance to *O. surinamensis*

In order to test whether the observed symbiont-mediated changes in cuticular thickness, melanization, and CHC composition affect desiccation resistance in *O. surinamensis*, we measured water loss and mortality of symbiotic and aposymbiotic beetles under desiccation stress. Indeed, aposymbiotic beetles reared at high or low humidity desiccated faster than their symbiotic counterparts (Table 1, Fig. 5a; measured as proportional decrease in weight as beetle dry mass differed between treatments; see Supplemental Fig. 5). Concordantly, symbiont-free beetles also exhibited higher mortality upon acute drought stress, independent of the humidity they experienced during development (Cox Mixed-Effect Model, N=400 (8 replicates with 50 individuals per treatment), Table 1, Fig. 5b). Similarly, survival from oviposition until emergence of adults was significantly lower in the absence of symbionts under low humidity (χ^2^ homogeneity test at 30% RH: 10.5% for aposymbiotic beetles, N=114, vs 27.5% for symbiotic beetles, N=40, χ^2^=6.71, p=0.013), but not at high humidity (χ^2^ homogeneity test at 60% RH: 31.4% for aposymbiotic beetles, N=296 vs 41.7% for symbiotic beetles, N=103, χ^2^=3.63, p=0.057). The differential susceptibility to desiccation was also reflected in the beetles’ population growth over three months, with a significant influence of symbiont absence, ambient humidity as well as their interaction (Table 1, Fig. 6).

**Figure 5.**
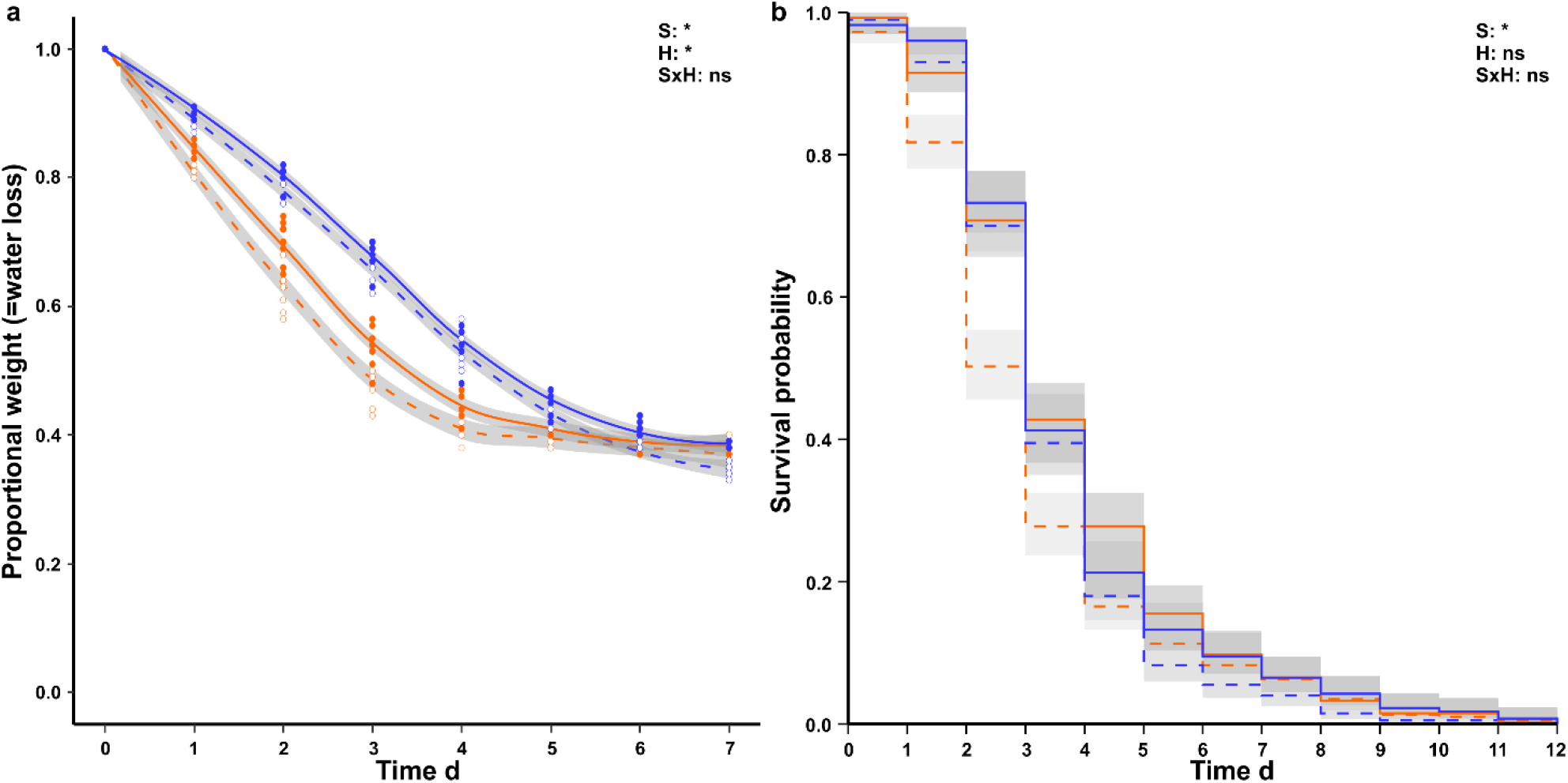
Water loss and survival of *O. surinamensis* adults under acute desiccation stress. (a) Water loss is significantly influenced by symbiont presence as well as rearing humidity (S,H; GLM, p<0.05), but not their interaction (S*H; GLM, p>0.05), whereas (b) mortality is only significantly influenced by symbiont presence (S; Cox Mixed-Effect Model, p<0.05). Lines show mean, and shaded areas 99% confidence intervals of (a) 50 pooled beetles for eight replicate populations per treatment and (b) 50 individual beetles from eight replicate populations per treatment. Continuous lines represent symbiotic and dashed lines aposymbiotic beetles, orange lines indicate rearing at 60% RH, blue lines at 30% RH

**Figure 6.**
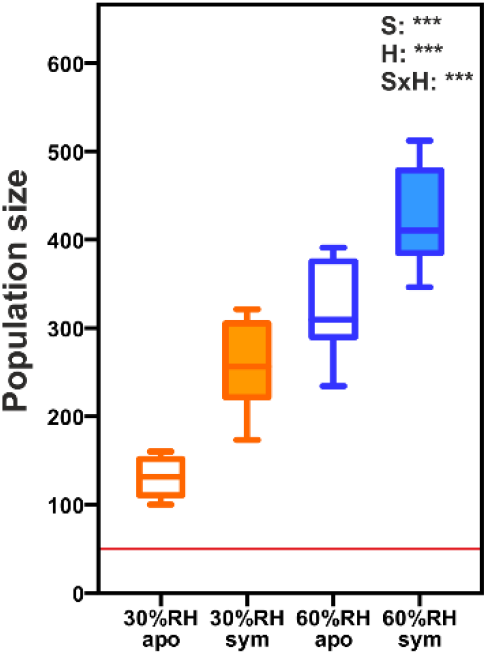
Population growth of *O.surinamensis* over three months from starting populations of 50 beetles. Symbiont presence, environmental humidity and their interaction have a significant influence on population growth (S, H, S*H; GLM, p<0.001). The red line indicates the initial population size. Boxplots show median, quartiles, and minimum/maximum of eight replicate populations per treatment. Filled boxes represent symbiotic and empty ones aposymbiotic beetles, orange boxes indicate rearing at 30% RH, blue ones at 60% RH

## Discussion

We showed that grain pest beetles in the beetle families Bostrichidae and Silvanidae engage in a symbiotic association with a group of Bacteroidetes bacteria that is closely related to *Sulcia* and *Blattabacterium*, the obligate nutritional mutualists of cicadas and cockroaches, respectively. However, in contrast to these (Lo et al., 2003, Takiya et al., 2006), the beetle symbiont 16S rRNA sequences revealed multiple acquisition, exchange and/or loss events, and the bacterial partners exhibited higher sequence divergences, indicating older associations and/or faster evolutionary rates than the *Sulcia* and *Blattabacterium* associations (Silva and Santos-Garcia, 2015). The common ancestor of *Blattabacterium, Sulcia* and the symbionts of the bostrichid and silvanid beetles was estimated to have lived around 394 Mya, predating the evolution of the beetles (Coleoptera) (Hunt et al., 2007). Considering that the common ancestor of Aucchenorrhyncha, Blattodea, Silvanidae and Bostrichidae existed around 380 to 390 Mya and gave rise to all holometabolous and most hemimetabolous insects (Misof et al., 2014), an ancient infection with the Bacteriodetes symbiont and subsequent losses in all but these four taxa seems to be an unlikely scenario. The more parsimonious explanation, especially considering the estimated age of the Bostrichidae (~150Mya) and Silvanidae (~180Mya; Hunt et al. 2007) involves at least six independent acquisition events (one in Aucchenorrhyncha, one in Blattodea, two in Silvanidae, and two in Bostrichidae) (Fig. 1 and Supplemental Fig. 1) and is reminiscent of the repeated acquisition of symbionts from a few clades of intracellular gamma-proteobacteria across diverse insect taxa (Husnik et al., 2011). Possibly, particular clades of bacteria were adapted to an insect-associated (possibly parasitic) lifestyle and - akin to extant *Wolbachia* infections - successfully spread through mixed vertical and horizontal transmission, but only those associations evolving towards mutual benefits proved to be stable over evolutionary timescales, resulting in the patchy distribution we observe today. Supporting this hypothesis, a basal clade of the bacteroidetes endosymbionts actually consists of three species that are described as male-killing endosymbionts in different ladybug beetles (Fig.1; Hurst et al., 1997, Hurst et al., 1999).

Within the bostrichid-Bacteroidetes association, we discovered three cases of multipartite symbioses with two different strains of the same bacterial clade, located in different bacteriome organs or compartments. While co-obligate symbionts have been described repeatedly across sap-feeding Hemiptera, they are usually co-localized in adjacent bacteriocytes to facilitate exchange of metabolic products or intermediates for jointly synthesized products (McCutcheon and von Dohlen, 2011, Wu et al., 2006). Alternatively, they are intermixed in the same bacteriocytes, as is the case in the fragmented *Hodgkinia* genomes in several cicada species (Van Leuven et al., 2014, Campbell et al., 2015). Interestingly, some of the Bostrichid species seem to have lost their ancestral symbiont, a theory already formulated by Huger (1956) who also occasionally observed the formation of additional, uninfected bacteriomes in *R. dominica*. This raises not only the question of the individual contribution of the single strains, but also the physiological and especially ecological consequence of symbiont acquisition, replacement or loss events for the respective beetle groups (Joy, 2013, Sudakaran et al., 2015, Sudakaran et al., 2017). Ancestrally, both groups of beetles inhabit rather humid environments. While Bostrichid beetles are described to inhabit and feed on sapwood, dying or dead trees, Silvanid beetles feed presumably on fungal detritivores of decomposing wood (Hunt et al., 2007, Thomas and Leschen, 2009). Finally, while the ecological niches of the investigated beetles are all characterized by low humidity, they differ widely in the available nutrient composition of their diet. Dry wood inhabited by the genus *Lyctus* probably represents the resource that is poorest in nitrogen (Hoadley, 1998); dried fruit and roots as preferred by *Dinoderus spp*. may be similarly unbalanced (Nations, 1990), whereas the germ tissue of different grains may contain sufficient nitrogen sources (Souci et al., 2009), which could have contributed to the loss of the basal symbiont lineage in those beetles. The insect cuticle is in general composed of chitin fibrils - a polymer of N-acetyl-glucosamine, which is synthesized from glucose, glutamine and acetyl-coenzyme A (Muthukrishnan et al., 2012) - in a complex with proteins (Hackman, 1974). The outer layer of the cuticle, the exocuticle, can be melanized and sclerotized and thereby becomes harder, darker and supposedly more water proof (Hackman, 1974, Gibbs and Rajpurohit, 2010). The fact, that, unlike in *S. oryzae* (Vigneron et al., 2014), the thickness of both endo- and exocuticle of *O. surinamensis* is reduced in the absence of the symbiont, suggests a more general contribution of nutrients than specifically Dopa or a similar precursor for cuticle melanization and sclerotization (Klein et al., 2016). Thus, nutritional benefits provided by Bacteriodetes symbionts might likely reach beyond individual amino acids or their precursors for cuticle melanization as in *S. oryzae* (Vigneron et al., 2014) to nitrogen recycling and essential amino acid and vitamin provisioning, as in cockroaches and the Auchenorrhyncha (Sabree et al., 2009, McCutcheon and Moran, 2010) or in carpenter ants (Gil et al., 2003, Degnan et al., 2005) and *Nardonella* harboring weevils (Kuriwada et al., 2010, Hosokawa et al., 2015). The individual contributions and interactions between the symbiont strains are interesting topics for future studies, especially given the facultative nature of the intracellular symbiosis and possibility for experimental manipulation of symbiont infection status.

In addition to identifying an ancient group of symbiotic *Sulcia*-related bacteria in diverse grain and wood pest beetles, we demonstrated an ecological benefit in the saw-toothed grain beetle *O. surinamensis* conferred by their non-obligate, yet prevalent, intracellular symbiont. By supporting cuticle synthesis, the bacteria confer desiccation resistance to their host, which constitutes a significant fitness benefit for the beetles, particularly under the dry conditions that characterize their anthropogenic habitat of granaries and other storage facilities (Hagstrum et al., 1996). Interestingly, grain weevils that occupy an almost identical ecological niche evolved a symbiotic association with γ-proteobacteria *(Sodalis pierantonius)* that supports cuticle biosynthesis in a similar manner (Heddi et al., 1999, Vigneron et al., 2014) and may hence contribute to drought tolerance. Likewise, carpenter ants of the genus *Camponotus* and the invasive ant *Cardiocondyla obscurior* evolved symbioses with the γ-proteobacteria *Blochmannia* and *Candidatus* Westeberhardia cardiocondylae, respectively, that support cuticle melanization through the synthesis of essential amino acids or tyrosine precursors, respectively (*Blochmannia*, de Souza et al., 2011, *Candidatus* Westeberhardia, Klein et al., 2016). However, in which way these ants benefit from enhanced cuticle melanization remains unclear. Ants as well as beetles my generally benefit from a thicker and harder cuticle as a first line of defense against many natural enemies, like predators, parasitoids and pathogens. For the grain beetles, we hypothesize that the symbioses with Bacteroidetes and γ-proteobacteria, respectively, constituted (pre)adaptations that enabled the three families to independently invade their niches of originally individual items of dried grain, fruit and wood, but proved to be especially advantageous to invade recent, anthropogenic stores of dry grain and fruit, as well as seasoned wood.

## Materials & Methods

### Insect cultures

*O. surinamensis, A. advena, R. dominica, P. truncatus, D. bifoveolatus*, and *D. porcellus* insect cultures were obtained from the Julius-Kühn-Institute / Federal Research Centre for Cultivated Plants (Berlin, Germany), *O. mercator* from Fera Science Ltd (Darlington, UK) and *L. brunneus* from the Federal Institute for Material Research and Testing (Berlin, Germany). Continuous symbiotic and aposymbiotic (see below) *O. surinamensis* cultures were maintained in 1.8L plastic containers, filled with 50g oat flakes, at 30°C and ambient humidity between 40% and 60% in the dark.

### DNA extraction, 16s rDNA cloning and sequencing

Total DNA was isolated individually from 25-30 adults per species using the Epicentre MasterPure^™^ Complete DNA and RNA Purification Kit (Illumina Inc., Madison, Winsconsin, USA).

Total 16s rDNA was amplified from 5-12 DNA extracts per species using universal eubacterial primers fD1 and rP2 (Welsburg et al. 1991). Reaction mixtures for PCR amplification consisted of 6.4μl distilled water, 1.25μl PCR Buffer, 0.25μl MgCl2, 1.5μl dNTPs, 1μl of each primer (each 10 pmol/μl), 0.1μl Taq polymerase (5U/μl) and 1μl template in a final volume of 12.5μL. The PCR temperature profile was 95°C for 3 minutes, followed by 30 cycles of 95°C, 60°C, and 72°C for 40s each, and a subsequent elongation step at 72°C for 10min. PCR products were purified with the Analytik Jena innuPREP PCRpure Kit (Jena, Germany).

Bacterial 16S rRNA amplicons from five individuals per beetle species were cloned with a pSC-A-amp/kan vector (Strata Clone PCR Cloning Kit, Agilent Technologies, Santa Clara, California, USA) into *Escherichia coli* K12. Vector insertion sequences of successfully transformed colonies were amplified by another PCR using the flanking primer pair M13_fwd and M13_rev. The PCR parameters and purification were identical as described above, except that an annealing temperature of 65°C was used, and entire cells from clone colonies were added to the PCR reaction mix as template. Bidirectional Sanger sequencing was performed in house to obtain the full sequence of the amplified 16S fragments using the M13_fwd/rev primers.

### Infection frequencies

Diagnostic PCRs were conducted to assess symbiont infection frequencies in all eight insect species. Specific oligonucleotides (Supplemental Table 1) were designed based on an alignment of *Sulcia muelleri, Blattabacterium sp*. and free-living Bacteriodetes 16S rDNA sequences. Specificity of the primer sequences for Bacteriodetes was assessed *in silico* using the Ribosomal Database Project 16S rDNA collection (Cole et al., 2014). Specificity was further tested by trying to amplify fragments from DNA extracts of the beetle species that should not carry the focal symbiont as well as European firebug *Pyrrhocoris apterus* and European beewolf *Philanthus triangulum* guts DNA extracts that similarly are known to lack Bacteroidetes bacteria.

### Phylogenetic inference

Beetle symbiont 16S rDNA sequences were aligned to representative *Sulcia muelleri, Blattabacterium* and free living Bacteriodetes 16S rDNA sequences obtained from the NCBI database, using the SILVA algorithm (Quast et al., 2013, Yilmaz et al., 2014). A maximum-likelihood phylogenetic tree was reconstructed with PHYML (Guindon and Gascuel, 2003) as implemented in Geneious Pro 9.1.5 (Drummond AJ, 2011), using the GTR model with uniform substitution rates per site. The initial tree for the heuristic search was obtained automatically by applying Neighbour-Joining and BioNJ algorithms to a matrix of pairwise distances estimated using the Maximum Composite Likelihood (MCL) approach, and then selecting the topology with superior log likelihood value. Bootstrap values were obtained from 10,000 replicates. A second tree was reconstructed by Bayesian inference applying a GTR +G +I model using MrBayes 3.2 (Huelsenbeck et al., 2001, Huelsenbeck and Ronquist, 2001, Ronquist and Huelsenbeck, 2003). The analysis ran for 2,000,000 generations, with trees sampled every 1,000 generations. After confirming that split frequencies converged to less than 0.01, we used a “Bumin” of 20%. Both trees were visualized with FigTree (http://tree.bio.ed.ac.uk/software/figtree/).

### Phylogenetic dating

Divergence time estimations were inferred using BEAST v1.8.4. MCMC analyses (Drummond A. J. and Rambaut, 2007) with HKY, GTR and TN93 nucleotide substitution models (empirical or estimated base frequencies, various site heterogeneity models [none, G, I+G]). Analyses were conducted under a strict clock (using a single rate of sequence evolution across the phylogeny) and an uncorrelated lognormal relaxed clock model (allowing variable substitution rates; Drummond A. J. et al., 2006). In each analysis, 100 million steps were performed, and trees were sampled every 100,000 steps. The phylogenetic tree from the Bayesian inference analysis (see previous section) was used a fixed input tree in all analyses. Nucleotide substitution priors were determined with jmodeltest v2.1.9 (Guindon and Gascuel, 2003, Darriba et al., 2012).

The age of the *cockroach-Blattabacterium* (normal distribution, mean±SD=220±25; Patino-Navarrete et al., 2013) and cicada-*Sulcia muelleri* (normal distribution, mean±=270±3.5; Moran et al., 2005) symbioses were used as calibration points for the dating analyses.

Selection of best fitting model was performed by path and stepping stone sampling (Baele and Lemey, 2013) with 100 steps logged every 1 million generations. The analysis with the best fitting model (GTR+I+G) was repeated twice with only one of both calibration points. Models were evaluated using Tracer v1.6 (http://beast.bio.ed.ac.uk/Tracer), consensus trees were generated with TreeAnnotator v1.8.4 (Drummond A. J. and Rambaut, 2007) using a burnin of 10% and a posterior probability limit of 0.3 and visualized with FigTree v1.4.2 (http://tree.bio.ed.ac.uk/software/figtree/).

### Fluorescence *in situ* hybridization (FISH)

Whole mount FISH was performed on *O.surinamensis* larvae, standard FISH on squashed, fixed eggs and on sections of adult beetles. Fresh or frozen beetle eggs were fixed by slightly squashing them on a glass slide and incubation for three minutes in 70% ethanol and another 3 minutes in 96% ethanol. Whole larvae, pupae and adults were briefly washed in diethylether and fixed for at least three days in 4% paraformaldehyde in PBS. Adults were then dehydrated and embedded in Technovit 8100 (Heraeus-Kulzer, Germany), and 10μm sections were cut on a microtome Microtome (Microm HM355S, Leica, Germany) and mounted on diagnostic microscope slides.

Probes were designed based on specific primers (see Supplemental Table 1) and were labelled with the cyanine dyes Cy3 or Cy5. Sections of adults and whole eggs were covered with hybridization buffer containing 0.9M NaCl, 0.02M Tris/HCl, 0.01% SDS, 0.5μM of each labelled oligonucleotide probe and 5μg/ml of the general DNA stain DAPI. Hybridization was performed at 50°C for 60min. The samples were subsequently washed twice with washing buffer consisting of 0.1M NaCl, 0.02M Tris/HCl, 5,mM EDTA and 0.01% SDS and incubated at 50°C for 20min in washing buffer, followed by a washing step with distilled water. After drying, the sections or eggs were covered with Vectashield (Vector Laboratories, Curlingham, CA, USA) and a cover slip.

Whole mount fish was performed by staining entire larvae at 50°C overnight in the same hybridization buffer. Afterwards, samples were washed twice for two hours with pre-warmed washing buffer at 50°C and twice for 20min with distilled water at room temperature, before mounting on 2-well slides and covering with Vectarshield for fluorescence microscopy.). Images were acquired with an AxioImager Z2 equipped with an Apotome.2 (Zeiss, Germany) and a SOLA light engine LED light source (Lumencor, OR, USA) under 200-400x magnification with the Z-stack option.

### Elimination of *O.surinamensis* symbionts

In order to obtain symbiont-free *O. surinamensis*, 150 adults were kept for three months on oat flakes that were soaked in a tetracycline solution (30mg tetracycline hydrochloride / g oat flakes; Sigma-Aldrich, Germany) and dried at 60°C. 200 adult offspring individuals of these beetles were then transferred back to a standard oat flake diet. A control group experienced the same conditions except that the tetracycline was omitted from the oat flake soaking step. Efficiency of symbiont elimination was verified by qPCR of both eggs and adults immediately after the tetracycline treatment as well as three and twelve months later, respectively. For all time points, eggs were also collected and squashed onto slides, fixed for 10min with 95% and 70% ethanol subjected to FISH as described above to verify symbiont absence. Furthermore, eight female adults of the F2-F3 generation (three months after the tetracycline treatment) were sectioned and subjected to FISH as described above.

Quantitative PCRs were carried out in 25μL reactions using the Qiagen QuantiTect-SYBR-Green-PCR mix (Qiagen, Venlo, The Netherlands), including 0.5μM of each primer and 1μL template DNA. To compensate for host developmental stage and size as well as DNA extraction efficiency, all qPCR samples were additionally subjected to a PCR with primers targeting the host 28S rRNA gene, and the resulting delta Ct values were used for relative quantification of bacterial 16S rRNA copies per host 28S rRNA copy (Pfaffl, 2001).

### Physiological response and fitness of symbiotic and aposymbiotic *O. surinamensis* lines

Eight symbiotic and aposymbiotic *O. surinamensis* populations were founded from one year old aposymbiotic and symbiotic control cultures and reared at 30-40% RH and 60% RH in a full-factorial design to measure cuticle thickness of 4^th^ instar larvae, melanization and thickness of the adult cuticle, cuticular hydrocarbon profiles of adults, desiccation resistance measured as water loss, as well as survival and population growth. For each replicate, 50 beetles were transferred to a box with oat flakes (20g) that were pre-conditioned for one week to the experimental conditions. Replicate populations were kept at 28°C and the two different humidity conditions in the dark for three months. In parallel to the replicate treatments, individual females from the basic cultures of all four treatments were separated into 12 well plates, eggs collected and the offspring individually reared in 48-well plates provided with one oat flake and incubated under above mentioned conditions. Survival until emergence of adults was monitored daily to assess mortality during development.

To evaluate the impact of humidity and symbiont elimination on cuticle melanisation, we determined the inverse red values (Vigneron et al., 2014) of 24 beetles from each treatment group. Photographs were taken with a Sony NEX 5 camera coupled to a Motic dissection stereoscope (Wetzlar, Germany) under identical conditions. Average red values were measured within an elliptic area covering the ventral thorax with the histogram tool in ImageJ 1.50a (Rasband, 1997-2016) and transformed into the inverse red values. To measure cuticle thickness, 9-14 adult beetles and ten 4^th^ instar larvae per treatment group were fixated in 4% paraformaldehyde in PBS. These beetles, as well as larvae from collected after three months from the eight replicate populations were embedded in epoxy resin (Epon_812 substitute, Sigma-Aldrich, Germany) and 1μm cross sections of the thorax next to the second pair of legs were cut on a Microtome (Microm HM355S, Leica, Germany) with a diamond blade and mounted on silanized glass slides with Histokitt (Roth, Germany). Images to measure cuticle diameter were taken with an AxioImager Z2 (Zeiss, Germany) under 200x magnification and differential interference contrast. Mean cuticle diameter was measured for one randomly chosen dorsal, ventral and lateral point, respectively, with the ZEN software distance tool (Zeiss, Germany).

Living adult beetles of each population were counted manually after three months to measure population growth. Immediately after counting, two batches of 50 beetles of each population were transferred to separate containers that were either empty or provided with three dried oat flakes, to measure water loss and survival, respectively. At the same time, two samples of 30 beetles each were transferred to 1.4mL glass vials to measure cuticular hydrocarbon profiles before and after desiccation stress, respectively.

Desiccation resistance was measured as water loss and survival rates by incubating containers of 50 beetles in a chamber that was covered with a layer of silica gel. The chamber was aerated with a constant air stream of 1 mL/min that was guided through a column of silica gel to reduce it’s humidity. The humidity inside the chamber was thereby reduced to below 2% RH within 30min after closing the box. One group of containers with 50 beetles was weighed daily to measure water loss keeping dead beetles in the container. From the other group, dead beetles were counted and removed daily to monitor survival.

To assess the impact of low humidity, symbiont elimination, and acute desiccation stress on CHC profiles, glass vials containing 30 symbiotic or aposymbiotic beetles that had been reared under high (60% RH) or low humidity (30-40% RH) were incubated for 24h in a desiccation chamber as described above and subsequently given another 24h to recover under their respective rearing conditions. Control groups were kept for 48h under the respective rearing conditions. Afterwards, all beetles from each vial were freeze-killed and extracted for 10 min with 100μL hexane HPLC-grade (Roth, Germany) containing 2μg octadecane (Sigma-Aldrich, Germany) as internal standard. After removal of beetles, extracts were concentrated to ~30μL, and 5μL were analysed on a Varian 450GC gas chromatograph coupled to a Varian 240MS ion-trap mass spectrometer (Agilent Technologies, Böblingen, Germany) using a split/splitless injector at 250°C with the purge valve opened after 60s. The GC was equipped with a DB5-ms column (30 m×0.25 mm ID; 0.25 μm df; Agilent, Santa Clara, CA, USA) and programmed from 150 to 320°C at 5°C/min with a 5 min. final isothermal hold. Helium was used as carrier gas, with a constant flow rate of 1ml/min. Mass spectra were recorded using electron ionization (EI-MS) with an ion trap temperature of 90°C. Data acquisition and quantifications were achieved with MS Workstation Version 6.9.3 Software (Agilent Technologies). Hydrocarbons were identified by retention index and fragmentation pattern in accordance with Howard et al. (1995). CHCs were automatically quantified using the Varian MS Workstation 6.9.3 software with manual correction. For analysis, we calculated total amount of CHCs per beetle (based on the amount of internal standard), the proportion of saturated CHCs, and a carbon chain length index (sum of the proportions of compounds multiplied by their respective number of carbon atoms).

### Statistical procedures

Symbiont abundance (ΔC_T_ (symbiont 16s rDNA/host 28S rDNA)), initial test of cuticle melanization and population growth were tested between treatment and control by exact, 2-sided Mann-Whitney-U tests in SPSS 23 (IBM, Armonk, NY).

Influence of symbiont infection, rearing humidity, and in case of CHC profile also desiccation stress, and their interaction effects was tested with generalized linear models (GLMs) in SPSS 23. For beetle melanization, cuticle diameter of adults and larvae, all CHC measurements and water loss, we used linear scale response models with a normal distribution. For the population size counts, we used a Poisson distribution and a log link function. Model parameters were estimated by the quasi-likelihood method and accepted if the full factorial model showed a significantly better fit than the intercept-only model (in all cases p<0.001). Wald χ^2^ statistics were calculated for the models and single factors. Boxplots were also visualized with SPSS 23. Water loss over time and across treatments was tested with generalized linear models with repeated measures, also in SPSS23 and visualized using the ‘ggplot2’ (Wickham and Chang, 2016) package in R studio version 3.1.1.

Mortality of *O.surinamensis* adults was analysed using a Cox mixed effects model with symbiont infection and rearing humidity and a random intercept per replicate population. The analysis was carried out using the package ‘coxme’ (Therneau, 2012) in R studio version 3.1.1. Survival probability of treatments was plotted based on Kaplan-Meier models using the ‘rms’ package (Harrell and Frank, 2013).

Survial during juvenile development (measured as percentage of individuals successfully developing from eggs to adults) was compared by manually calculating χ^2^ homogeneity tests.

## Data availability

Partial symbiont 16S rDNA sequences are deposited in Genbank under accession numbers MF183956-MF183966.

## Contributions

T.E. and M.K. designed the project and wrote the manuscript, C.A. and R.P. provided beetle cultures and specimen, N.E., C.G. and T.K. characterized the symbionts of *O. surinamensis*, T.S. performed *O.surinamensis* larval survival assays, T.E. performed all other experiments and analyzed the data.

## Acknowledgements

M.K. and T.E. acknowledge funding from the Max-Planck-Society. We thank Benjamin Weiss, Dagmar Klebsch and Christiane Stürzbecher for technical assistance.

## References (50 + unlimited in Methods only)

Amann RI, Binder BJ, Olson RJ, Chisholm SW, Devereux R & Stahl DA. 1990. Combination of 16s rRNA-Targeted Oligonucleotide Probes with Flow-Cytometry for Analyzing Mixed Microbial-Populations. Appl. Environ. Microbiol. 56: 1919–1925.

Baele G & Lemey P. 2013. Bayesian evolutionary model testing in the phylogenomics era: matching model complexity with computational efficiency. Bioinformatics 29: 1970–1979, doi:10.1093/bioinformatics/btt340.

Brumin M, Kontsedalov S & Ghanim M. 2011. *Rickettsia* influences thermotolerance in the whitefly *Bemisia tabaci* B biotype. Insect Sci. 18: 57–66, doi:10.1111/j.1744-7917.2010.01396.x.

Buchner P. 1965. Endosymbiosis of Animals with Plant Microorganisms. New York: John Wiley & Sons

Campbell MA, Van Leuven JT, Meister RC, Carey KM, Simon C & Mccutcheon JP. 2015. Genome expansion via lineage splitting and genome reduction in the cicada endosymbiont Hodgkinia. Proc. Natl. Acad. Sci. U. S. A. 112: 10192–10199, doi:10.1073/pnas.1421386112.

Cole JR, Wang Q, Fish JA, Chai BL, Mcgarrell DM, Sun YN, Brown CT, Porras-Alfaro A, Kuske CR & Tiedje JM. 2014. Ribosomal Database Project: data and tools for high throughput rRNA analysis. Nucleic Acids Res. 42: D633–D642, doi:10.1093/nar/gkt1244.

Corbin C, Heyworth ER, Ferrari J & Hurst GDD. 2017. Heritable symbionts in a world of varying temperature. Heredity 118: 10–20, doi:10.1038/hdy.2016.71.

Crowson RA. 1981. The Biology of the Coleoptera. London, UK: Academic Press

Darriba D, Taboada GL, Doallo R & Posada D. 2012. jModelTest 2: more models, new heuristics and parallel computing. Nat. Meth. 9: 772–772.

De Souza JD, Devers S & Lenoir A. 2011. *Blochmannia* endosymbionts and their host, the ant *Camponotus fellah*: Cuticular hydrocarbons and melanization. C. R. Biol. 334: 737–741, doi:10.1016/j.crvi.2011.06.008.

Degnan PH, Lazarus AB & Wernegreen JJ. 2005. Genome sequence of *Blochmannia pennsylvanicus* indicates parallel evolutionary trends among bacterial mutualists of insects. Genome Res. 15: 1023–1033, doi:10.1101/Gr.3771305.

Douglas AE. 2009. The microbial dimension in insect nutritional ecology. Funct. Ecol. 23: 38–47, doi:10.1111/j.1365-2435.2008.01442.x.

Drummond A. 2011. Geneious v9.1.5. Auckland, New Zealand: Biomatters Ltd.

Drummond AJ, Ho SYW, Phillips MJ & Rambaut A. 2006. Relaxed phylogenetics and dating with confidence. PLoS Biol. 4: 699–710, doi:10.1371/journal.pbio.0040088.

Drummond AJ & Rambaut A. 2007. BEAST: Bayesian evolutionary analysis by sampling trees. BMCEvol. Biol. 7, doi:10.1186/1471-2148-7-214.

Feldhaar H. 2011. Bacterial symbionts as mediators of ecologically important traits of insect hosts. Ecol. Entomol. 36: 533–543, doi:10.1111/j.1365-2311.2011.01318.x.

Florez LV, Biedermann PHW, Engl T & Kaltenpoth M. 2015. Defensive symbioses of animals with prokaryotic and eukaryotic microorganisms. Nat. Prod. Rep. 32: 904–936, doi:10.1039/C5NP00010F.

Gibbs AG & Rajpurohit S. 2010. Cuticular Lipids and Water Balance. In: Bagnères A-G & Blomquist GJ, editors. Insect Hydrocarbons: Biology, Biochemistry, and Chemical Ecology. Cambridge: Cambridge University Press. p100–120

Gil R, Silva FJ, Zientz E, Delmotte F, Gonzalez-Candelas F, Latorre A, Rausell C, Kamerbeek J, Gadau J, Holldobler B, Van Ham RCHJ, Gross R & Moya A. 2003. The genome sequence of *Blochmannia floridanus*: Comparative analysis of reduced genomes. Proc. Natl. Acad. Sci. U. S. A. 100: 9388–9393, doi:10.1073/pnas.1533499100.

Guindon S & Gascuel O. 2003. A simple, fast, and accurate algorithm to estimate large phylogenies by maximum likelihood. Syst. Biol. 52: 696–704, doi:10.1080/10635150390235520.

Hackman. 1974. Chemistry of the insect cuticle. In: Rockenstein M, editor The Physiology Of Insecta. New York, NY, USA: Academic Press. p215–270

Hagstrum DW, Flinn PW & Howard RW. 1996. Ecology. In: Hagstrum DW & Subramanyam B, editors. Integrated Management of Insects in Stored Products. New York, NY, USA: Marcel Dekker. p71–134

Harmon JP, Moran NA & Ives AR. 2009. Species Response to Environmental Change: Impacts of Food Web Interactions and Evolution. Science 323: 1347–1350, doi:10.1126/science.1167396.

Harrell J & Frank E. 2013. Rrms: Regression Modeling Strategies. R package version 5.1-0

Heddi A, Grenier AM, Khatchadourian C, Charles H & Nardon P. 1999. Four intracellular genomes direct weevil biology: Nuclear, mitochondrial, principal endosymbiont, and Wolbachia. Proc. Natl. Acad. Sci. U. S. A. 96: 6814–6819, doi:10.1073/pnas.96.12.6814.

Hoadley RB. 1998. Chemical and Physical Properties of Wood. In: Dardes K & Rothe A, editors. The Structural Conservation of Panel Paintings. Los Angeles, CA, USA: Getty Publications.

Hosokawa T, Koga R, Tanaka K, Moriyama M, Anbutsu H & Fukatsu T. 2015. *Nardonella* endosymbionts of Japanese pest and non-pest weevils (Coleoptera: Curculionidae). Appl. Entomol. Zool. 50: 223–229, doi:10.1007/s13355-015-0326-y.

Howard RW, Howard CD & Colquhoun S. 1995. Ontogenic and Environmentally-Induced Changes in Cuticular Hydrocarbons of *Oryzaephilus surinamensis* (Coleoptera, Cucujidae). Ann. Entomol. Soc. Am. 88: 485–495.

Huelsenbeck JP & Ronquist F. 2001. MRBAYES: Bayesian inference of phylogenetic trees. Bioinformatics 17: 754–755, doi:10.1093/bioinformatics/17.8.754.

Huelsenbeck JP, Ronquist F, Nielsen R & Bollback JP. 2001. Evolution - Bayesian inference of phylogeny and its impact on evolutionary biology. Science 294: 2310–2314, doi:10.1126/science.1065889.

Huger A. 1956. Experimentelle Untersuchungen über die künstliche Symbiontenelimination bei Vorratsschädlingen: *Rhizopertha dominica* F. (Bostrychidae) und *Oryzaephilus surinamensis* L. (Cucujidae). Z. Morphol. Oekol. Tiere 44: 626–701, doi:10.1007/BF00390698.

Hunt T, Bergsten J, Levkanicova Z, Papadopoulou A, John OS, Wild R, Hammond PM, Ahrens D, Balke M, Caterino MS, Gómez-Zurita J, Ribera I, Barraclough TG, Bocakova M, Bocak L & Vogler AP. 2007. A Comprehensive Phylogeny of Beetles Reveals the Evolutionary Origins of a Superradiation. Science 318: 1913–1916, doi:10.1126/science.1146954.

Hurst GDD, Bandi C, Sacchi L, Cochrane AG, Bertrand D, Karaca I & Majerus MEN. 1999. *Adonia variegata* (Coleoptera: Coccinellidae) bears maternally inherited Flavobacteria that kill males only. Parasitology 118: 125–134, doi:10.1017/S0031182098003655.

Hurst GDD, Hammarton TC, Bandi C, Majerus TMO, Bertrand D & Majerus MEN. 1997. The diversity of inherited parasites of insects: the male-killing agent of the ladybird beetle *Coleomegilla maculata* is a member of the Flavobacteria. Genet. Res. 70: 1–6, doi:10.1017/S0016672397002838.

Husnik F, Chrudimsky T & Hypsa V. 2011. Multiple origins of endosymbiosis within the Enterobacteriaceae (gamma-Proteobacteria): convergence of complex phylogenetic approaches. BMC Biol. 9, doi:10.1186/1741-7007-9-87.

Joy JB. 2013. Symbiosis catalyses niche expansion and diversification. Proc. R. Soc. Lond. B Biol. Sci. 280, doi:10.1098/Rspb.2012.2820.

Kleespies RG, Nansen C, Adouhoun T & Huger AM. 2001. Ultrastructure of bacteriomes and their sensitivity to ambient temperatures in *Prostephanus truncatus* (Horn). Biocontrol Sci. Technol. 11: 217–232, doi:10.1080/09583150120035648.

Klein A, Schrader L, Gil R, Manzano-Marin A, Florez L, Wheeler D, Werren JH, Latorre A, Heinze J, Kaltenpoth M, Moya A & Oettler J. 2016. A novel intracellular mutualistic bacterium in the invasive ant Cardiocondyla obscurior. ISME J. 10: 376–388, doi:10.1038/ismej.2015.119.

Klepzig KD, Adams AS, Handelsman J & Raffa KF. 2009. Symbioses: A Key Driver of Insect Physiological Processes, Ecological Interactions, Evolutionary Diversification, and Impacts on Humans. Environ. Entomol. 38: 67–77, doi:10.1603/022.038.0109.

Koch A. 1931. Die Symbiose von Oryzaephilus surinamensis L. (Cucujidae, Coleoptera). Z. Morphol. Oekol. Tiere 23: 389–424, doi:10.1007/BF00446355.

Koch A. 1936a. Symbiosestudien. I. Die Symbiose des Splintkäfers, *Lyctus linearis* Goeze. Z. Morphol. Oekol. Tiere 32: 92–136, doi:10.1007/BF00406593.

Koch A. 1936b. Symbiosestudien. II. Experimentelle Untersuchungen an *Oryzaephilus surinamensis* L. (Cucujidae, Coleopt.). Z. Morphol. Oekol. Tiere 32: 137–180, doi:10.1007/BF00406594.

Koch A. 1956. The experimental elimination of symbionts and its consequences. Exp. Parasitol. 5: 481–518, doi:10.1016/S0014-4894(56)80008-8.

Kuriwada T, Hosokawa T, Kumano N, Shiromoto K, Haraguchi D & Fukatsu T. 2010. Biological Role of *Nardonella* Endosymbiont in Its Weevil Host. PLoS One 5, doi:10.1371/journal.pone.0013101.

Levkanicova Z. 2009. Molecular Phylogeny of the Superfamily Tenebrionoidea (Coleoptera: Cucujiformia). PhD, Palacky Univarsity.

Lo N, Bandi C, Watanabe H, Nalepa C & Beninati T. 2003. Evidence for cocladogenesis between diverse dictyopteran lineages and their intracellular endosymbionts. Mol. Biol. Evol. 20: 907–913, doi:10.1093/molbev/msg097.

Mansour K. 1934. On the Intracellular Micro-organisms of some Bostrychid Beetles. Q. J. Microsc. Sci. 77: 243–U17.

Mccutcheon JP & Moran NA. 2010. Functional Convergence in Reduced Genomes of Bacterial Symbionts Spanning 200 My of Evolution. Genome Biol. Evol. 2: 708–718, doi:10.1093/gbe/evq055.

Mccutcheon JP & Von Dohlen CD. 2011. An Interdependent Metabolic Patchwork in the Nested Symbiosis of Mealybugs. Curr. Biol. 21: 1366–1372, doi:10.1016/j.cub.2011.06.051.

Misof B, Liu SL, Meusemann K, Peters RS, Donath A, Mayer C, Frandsen PB, Ware J, Flouri T, Beutel RG, Niehuis O, Petersen M, Izquierdo-Carrasco F, Wappler T, Rust J, Aberer AJ, Aspock U, Aspock H, Bartel D, Blanke A, Berger S, Bohm A, Buckley TR, Calcott B, Chen JQ, Friedrich F, Fukui M, Fujita M, Greve C, Grobe P, Gu SC, Huang Y, Jermiin LS, Kawahara AY, Krogmann L, Kubiak M, Lanfear R, Letsch H, Li YY, Li ZY, Li JG, Lu HR, Machida R, Mashimo Y, Kapli P, Mckenna DD, Meng GL, Nakagaki Y, Navarrete-Heredia JL, Ott M, Ou YX, Pass G, Podsiadlowski L, Pohl H, Von Reumont BM, Schutte K, Sekiya K, Shimizu S, Slipinski A, Stamatakis A, Song WH, Su X, Szucsich NU, Tan MH, Tan XM, Tang M, Tang JB, Timelthaler G, Tomizuka S, Trautwein M, Tong XL, Uchifune T, Walzl MG, Wiegmann BM, Wilbrandt J, Wipfler B, Wong TKF, Wu Q, Wu GX, Xie YL, Yang SZ, Yang Q, Yeates DK, Yoshizawa K, Zhang Q, Zhang R, Zhang WW, Zhang YH, Zhao J, Zhou CR, Zhou LL, Ziesmann T, Zou SJ, Li YR, Xu X, Zhang Y, Yang HM, Wang J, Wang J, Kjer KM, et al. 2014. Phylogenomics resolves the timing and pattern of insect evolution. Science 346: 763–767, doi:10.1126/science.1257570.

Montllor CB, Maxmen A & Purcell AH. 2002. Facultative bacterial endosymbionts benefit pea aphids *Acyrthosiphon pisum* under heat stress. Ecol. Entomol. 27: 189–195, doi:10.1046/j.1365-2311.2002.00393.x.

Moran NA. 2006. Symbiosis. Curr. Biol. 16: R866–R871, doi:10.1016/j.cub.2006.09.019.

Moran NA, Tran P & Gerardo NM. 2005. Symbiosis and Insect Diversification: an Ancient Symbiont of Sap-Feeding Insects from the Bacterial Phylum Bacteroidetes. Appl. Environ. Microbiol. 71: 8802–8810, doi:10.1128/aem.71.12.8802-8810.2005.

Muthukrishnan S, Merzendorfer H, Arakane Y & Kramer KJ. 2012. Chitin Metabolism in Insects. In: Gilbert LI, editor Insect Molecular Biology and Biochemistry. San Diego, CA, USA: Academic Press. p193–235

Nardon P & Grenier AM 1988. Genetical and Biochemical Interactions Between the Host and its Endocytobiotes in the Weevils *Sitophilus* (Coleoptere, Curculionidae) and other related species. In: Scannerini S, Smith D, Bonfante-Fasolo P & Gianinazzi-Pearson V (eds.) Cell to Cell Signals in Plant, Animal and Microbial Symbiosis. Berlin, Germany: Springer-Verlag.

Nations FaaOOTU. 1990. Roots, tubers, plantains and bananas in human nutrition. food & Agriculture Organization

Oliver KM & Martinez AJ. 2014. How resident microbes modulate ecologically important traits of insects. Curr. Opin. Insect Sci. 4: 1–7, doi:10.1016/j.cois.2014.08.001.

Patino-Navarrete R, Moya A, Latorre A & Pereto J. 2013. Comparative Genomics of *Blattabacterium cuenoti:* The Frozen Legacy of an Ancient Endosymbiont Genome. Genome Biol. Evol. 5: 351–361, doi:10.1093/gbe/evt011.

Pfaffl MW. 2001. A new mathematical model for relative quantification in real-time RT-PCR. Nucleic Acids Res. 29, doi:10.1093/nar/29.9.e45.

Quast C, Pruesse E, Yilmaz P, Gerken J, Schweer T, Yarza P, Peplies J & Glockner FO. 2013. The SILVA ribosomal RNA gene database project: improved data processing and web-based tools. Nucleic Acids Res. 41: D590–D596, doi:10.1093/nar/gks1219.

Rasband WS. 1997-2016. ImageJ. Bethesda, Maryland, USA: U. S. National Institutes of Health

Ronquist F & Huelsenbeck JP. 2003. MrBayes 3: Bayesian phylogenetic inference under mixed models. Bioinformatics 19: 1572–1574, doi:10.1093/bioinformatics/btg180.

Russell JA & Moran NA. 2006. Costs and benefits of symbiont infection in aphids: variation among symbionts and across temperatures. Proc. R. Soc. Lond. B Biol. Sci. 273: 603–610, doi:10.1098/rspb.2005.3348.

Sabree ZL, Kambhampati S & Moran NA. 2009. Nitrogen recycling and nutritional provisioning by *Blattabacterium*, the cockroach endosymbiont. Proc. Natl. Acad. Sci. U. S. A. 106: 19521–19526, doi:10.1073/pnas.0907504106.

Silva FJ & Santos-Garcia D. 2015. Slow and Fast Evolving Endosymbiont Lineages: Positive Correlation between the Rates of Synonymous and Non-Synonymous Substitution. Front. Microbiol. 6, doi:10.3389/Fmicb.2015.01279.

Souci SW, Fachmann W & Kraut H. 2009. Lebensmitteltabelle für die Praxis. Stuttgart: Wissenschaftliche Verlagsgesellschaft

Sudakaran S, Kost C & Kaltenpoth M. 2017. Symbiont Acquisition and Replacement as a Source of Ecological Innovation. Trends Microbiol., doi:10.1016/j.tim.2017.02.014.

Sudakaran S, Retz F, Kikuchi Y, Kost C & Kaltenpoth M. 2015. Evolutionary transition in symbiotic syndromes enabled diversification of phytophagous insects on an imbalanced diet. ISME J. 9: 2587–2604, doi:10.1038/ismej.2015.75.

Takiya DM, Tran PL, Dietrich CH & Moran NA. 2006. Co-cladogenesis spanning three phyla: leafhoppers (Insecta: Hemiptera: Cicadellidae) and their dual bacterial symbionts. Mol. Ecol. 15: 4175–4191, doi:10.1111/j.1365-294X.2006.03071.x.

Therneau T. 2012. Coxme: Mixed Effects Cox Models. R package version 2.2-5

Thomas MC & Leschen RaB. 2009. Silvanidae Kirby, 1837. In: Leschen RaB, Beutel RG & Lawrence JF, editors. Handbook of Zoology - Coleoptera, Beetles. Berlin, Germany: De Gruyter. p346–350

Van Den Bosch TJM & Welte CU. 2017. Detoxifying symbionts in agriculturally important pest insects. Microb. Biotechnol. 10: 531–540, doi:10.1111/1751-7915.12483.

Van Leuven JT, Meister RC, Simon C & Mccutcheon JP. 2014. Sympatric Speciation in a Bacterial Endosymbiont Results in Two Genomes with the Functionality of One. Cell 158: 1270–1280, doi:10.1016/j.cell.2014.07.047.

Vigneron A, Masson F, Vallier A, Balmand S, Rey M, Vincent-Monegat C, Aksoy E, Aubailly-Giraud E, Zaidman-Remy A & Heddi A. 2014. Insects Recycle Endosymbionts when the Benefit Is Over. Curr. Biol. 24: 2267–2273, doi:10.1016/j.cub.2014.07.065.

Weisburg WG, Barns SM, Pelletier DA & Lane DJ. 1991. 16S Ribosomal DNA Amplification for Phylogenetic Study. J. Bacteriol. 173: 697–703, doi:10.1128/jb.173.2.697-703.1991.

Weller R, Glockner FO & Amann R. 2000. 16S rRNA-targeted oligonucleotide probes for the *in situ* detection of members of the phylum Cytophaga-Flavobacterim-Bacteroides. Syst. Appl. Microbiol. 23: 107114, doi:10.1016/S0723-2020(00)80051-X.

Wernegreen JJ. 2012. Mutualism meltdown in insects: bacteria constrain thermal adaptation. Curr. Opin. Microbiol. 15: 255–262, doi:10.1016/j.mib.2012.02.001.

Wickham H & Chang W. 2016. Create Elegant Data Visualisations Using the Grammar of Graphics. R package version 2.2.1

Wu D, Daugherty SC, Van Aken SE, Pai GH, Watkins KL, Khouri H, Tallon LJ, Zaborsky JM, Dunbar HE, Tran PL, Moran NA & Eisen JA. 2006. Metabolic Complementarity and Genomics of the Dual Bacterial Symbiosis of Sharpshooters. PLoS Biol. 4: e188, doi:10.1371/journal.pbio.0040188.

Yilmaz P, Parfrey LW, Yarza P, Gerken J, Pruesse E, Quast C, Schweer T, Peplies J, Ludwig W & Glockner FO. 2014. The SILVA and “All-species Living Tree Project (LTP)” taxonomic frameworks. Nucleic Acids Res. 42: D643–D648, doi:10.1093/nar/gkt1209.

